# Population collapse of the Egyptian fruit bat on Cyprus (2005−2022) was likely driven by roost disturbance and declining food availability linked to climate change

**DOI:** 10.64898/2026.04.09.717500

**Authors:** Radek K. Lučan, Haris Nicolaou, Tomáš Bartonička, Erik Bachorec, Martin Šálek, Šimon Řeřucha, Petr Jedlička, Elena Erotokritou, Ivan Horáček

## Abstract

Peripheral island populations are often especially vulnerable to environmental change, yet they may also represent unique components of biodiversity. We assessed long-term population change in the Egyptian fruit bat (Rousettus aegyptiacus) on Cyprus, the only insular and geographically isolated population of this pteropodid in Europe, and evaluated two non-exclusive explanations for its decline: roost disturbance and reduced food availability. We analysed roost counts from 21 underground sites monitored between 2005 and 2022 and modelled temporal trends in commercially produced fruits used by the species. The monitored population declined from c. 7200 to c. 1050 individuals, corresponding to an estimated decrease of 85.4%. The decline was steepest during 2005–2011, slowed during 2012–2017, and was followed by partial recovery in 2018–2022. Colonies in easily accessible roosts declined significantly faster than those in less accessible roosts, consistent with an important role of human disturbance. Fruit production showed strong long-term declines and multiple structural breaks clustered in the mid-2000s, coinciding with the most severe phase of population decline and a major drought period on Cyprus. The fate of the missing portion of the population remains uncertain. Although large-scale mortality cannot be excluded, there was no clear evidence of widespread starvation-related mortality, and emigration to nearby mainland areas remains a plausible but untested explanation. Overall, our results indicate that the collapse of this peripheral island population was most likely driven by a combination of roost disturbance and reduced food availability associated with climate-related environmental change, highlighting the urgent need for strict roost protection and measures to secure food and water resources.

## 1. Introduction

Populations occurring at the margins of species’ geographic ranges may be particularly vulnerable to environmental change. Peripheral populations often persist under environmental conditions that differ from those prevailing in the core of the distribution, which can lead to smaller population sizes, reduced connectivity, and increased demographic stochasticity (Brown, 1984; Lesica and Allendorf, 1995; Sexton et al., 2009). Consequently, such populations may be more prone to local extinction under changing environmental conditions (Channell and Lomolino, 2000; Eckert et al., 2008). At the same time, peripheral populations may harbour unique genetic variation or represent locally adapted lineages, thereby contributing disproportionately to the evolutionary potential and long-term persistence of species (Lesica and Allendorf, 1995; Hampe and Petit, 2005). These conservation concerns are even more pronounced in insular environments, where geographic isolation, limited immigration, and small population sizes increase susceptibility to demographic fluctuations and stochastic events (MacArthur and Wilson, 1967; Frankham, 1998; Whittaker and Fernández-Palacios, 2007). Populations located simultaneously at the edge of a species’ range and on islands therefore represent particularly vulnerable, yet often biologically unique, components of biodiversity.

Among bats, the Egyptian fruit bat (*Rousettus aegyptiacus*) population on Cyprus fits this definition particularly well. Eastern Mediterranean populations of this species represent the only extension of Old World fruit bats (Pteropodidae) beyond the tropics and form the northern margin of the species’ distribution range (Benda et al., 2011; Weinberg et al., 2023). Owing to the presence of a genetically isolated population on Cyprus (Hulva et al., 2012), the Egyptian fruit bat forms part of the European fauna and extends the list of mammalian families occurring in Europe to include Pteropodidae (Dietz et al., 2009).

Throughout much of its Mediterranean range, however, the species has traditionally been regarded as an agricultural pest and was subjected to extensive eradication campaigns during much of the 20th century (Korine et al. 1999). In Cyprus, direct killing of fruit bats, together with systematic destruction of their roosts, led to a substantial decline in the local population during the second half of the 20th century (Spitzenberger, 1979; Hadjisterkotis, 2006). In 1993, the official eradication campaign was halted, and the population appeared to stabilise or even recover during the following one to two decades (Hadjisterkotis, 2006). Accordingly, Benda et al. (2007) listed the Egyptian fruit bat among the most common bat species on the island based on the number of recent records. However, no detailed analysis of population trends has been published since Hadjisterkotis (2006).

The aims of this study were therefore to: (1) summarise all available data on population trends in the Cypriot population of the Egyptian fruit bat between 2005 and 2022; and (2) analyse possible causes of the observed decline, with particular attention to the potential roles of roost disturbance and decreasing food availability represented by cultivated fruits.

To assess the potential role of human pressure through roost disturbance, we hypothesised that colonies in easily accessible roosts should decline more strongly than those in less accessible roosts, because disturbance by people is more likely in the former. In evaluating the potential causes of reduced population viability, we considered the well-established fact that a substantial proportion of the Egyptian fruit bat’s diet consists of fruit species cultivated by humans (Korine et al., 1999; Del Vaglio et al., 2011; Aziz et al., 2016). Therefore, we hypothesised that major changes in fruit production, reflected in an accumulation of structural breaks in production dynamics, should coincide with or precede the period of the most pronounced population decline.

## 2. Materials and methods

### 2.1. Fruit bat roosts and monitoring

Using published sources, information provided by local people, and our own field surveys, we identified and inspected all underground spaces (both caves and mines) that could potentially serve as roosts for fruit bats. In total, we monitored 21 such sites (Fig. 1) annually in 14 of the 18 years between 2005 and 2022. Roost counts were conducted during the reproductive season (March–October) by direct visual inspection. Most roosts consisted of relatively small underground spaces, allowing straightforward visual estimates of the number of bats present. In roosts containing larger colonies, colony size was estimated from photographs taken during inspections. Most counts were rounded to the nearest 50 or 100 individuals.

**Figure 1.**
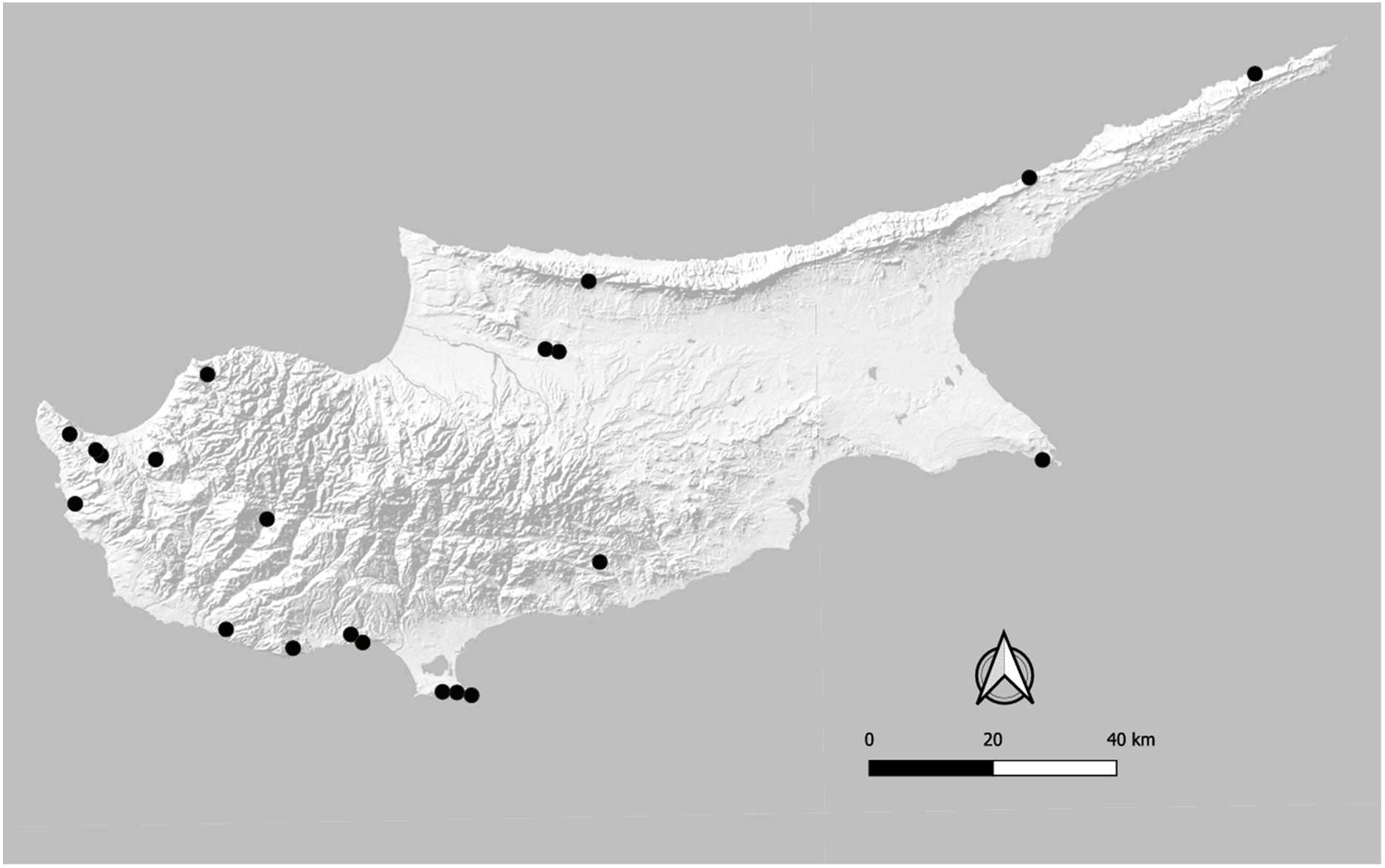
Map of Cyprus with marked locations (black dots) of fruitbat colonies monitored between 2005 and 2022.

To evaluate the possible effect of human disturbance in subsequent analyses, monitored roosts were classified into two categories according to their accessibility to people, which served as a proxy for the expected frequency of human roost disturbance. The two accessibility classes were defined as follows: (1) easy: roosts located close to human settlements, frequently used roads or tourist paths, often with an entrance visible from a long distance and without the need to climb steep rocky terrain or walk more than 100 m through dense thorny vegetation; (2) hard: roosts located far from frequently used roads or paths, with inconspicuous entrances and access requiring a long walk through difficult terrain, including dense thorny vegetation or rock climbing, or access by boat in the case of sea caves.

### 2.2. Commercial fruit production

Because the diet of the Egyptian fruit bat includes a wide range of fruits, many of which are commercially grown (Aziz et al., 2016), we compiled long-term data on fruit production in Cyprus from the Ministry of Agriculture, Rural Development and Environment of Cyprus, as reported through the Statistical Service of Cyprus to the Food and Agriculture Organization (FAO) and made available in the FAO database. We extracted annual production data for all fruit types known to be consumed by the Egyptian fruit bat (Aziz et al. 2016). These included apples (*Malus domestica*), apricots (*Prunus armeniaca*), bananas (*Musa* spp.), carob (*Ceratonia siliqua*), cherries (*Prunus cerasus*), figs (*Ficus carica*), grapes (*Vitis vinifera*), kiwi fruit (*Actinidia deliciosa*), oranges and other citrus fruits (*Citrus* spp.), peaches and nectarines (*Prunus persica*), pears (*Pyrus communis*), and plums and sloes (*Prunus domestica*).

### 2.4. Statistical analysis

#### 2.4.1. Population trends

Roost count data were reshaped into long format, with one record per roost–year combination and the following variables: site (roost identity), year (calendar year), count (observed abundance), and accessibility (roost accessibility category).

Long-term population trends were analysed using log-linear Poisson regression models implemented in the R package *rtrim* (Bogaart et al., 2025). In TRIM (Trends and Indices for Monitoring Data), model parameters are estimated by maximum likelihood within a Generalized Estimation Equations (GEE) framework, allowing correction for overdispersion and serial correlation in count data. Missing counts for sites not surveyed in particular years were estimated (“imputed”) from changes observed at all other sites. Counts were modelled with roost-specific effects and a time effect estimated by maximum likelihood. Calendar years were converted into a sequential time index corresponding only to monitored years. Because monitoring intervals were irregular, estimated slopes per TRIM time step were annualised by dividing by the mean interval (in years) between consecutive monitored calendar years. Annual percentage change and corresponding 95% confidence intervals were derived by transforming the annualised log-scale slope and its standard error using normal-approximation limits. Trend significance was assessed using Wald tests of the slope parameter.

To evaluate temporal heterogeneity in the population trajectory, we first fitted a TRIM time-effects model (model 3), which estimates a separate effect for each monitored year. Imputed annual indices and their standard errors were extracted and log-transformed. Year-to-year changes in log-indices were examined, and abrupt changes were assessed using Wald z-tests based on the differences between adjacent log-indices and their associated standard errors, assuming independence as a conservative approximation. This exploratory step was used to identify candidate breakpoints in the temporal trajectory.

Based on these results, the monitoring period was divided into three data-informed phases (2005–2011, 2012–2017, and 2018–2022). For each phase, a separate TRIM linear trend model was fitted, and annual percentage change was calculated as described above. Differences among period-specific slopes were evaluated using Wald z-tests comparing annualised slope estimates. To account for multiple pairwise comparisons among phases, p-values were adjusted using the Holm–Bonferroni method.

#### 2.4.2. Effect of roost accessibility on population trends

To test whether population trends differed between roost accessibility categories, we fitted a generalized linear mixed model in the R package *glmmTMB* (Brooks et al., 2017). Counts were modelled using a negative binomial error distribution (NB2 parameterisation) with a log link to account for overdispersion. Fixed effects included centred calendar year, roost accessibility category, and their interaction. Roost identity was included as a random intercept to account for repeated measurements and persistent differences in baseline abundance among sites. Observations with missing counts were excluded automatically during model fitting.

Accessibility-specific annual trends were derived from the fitted model as category-specific slopes of the year effect. For the reference category (easy), the slope corresponded to the main effect of year. The slope for hard-access roosts was obtained by adding the interaction coefficient (year × accessibility) to the reference slope. Standard errors for these derived slopes were calculated from the fixed-effect variance–covariance matrix, including covariance terms. Slopes were then transformed from the log scale to annual percentage change, and 95% confidence intervals were calculated using normal-approximation limits. Differences in trends between accessibility categories were evaluated using a Wald test of the interaction term (year × accessibility).

#### 2.4.3. Long-term changes in agricultural fruit production

Annual production data were compiled for the period 1990–2022 for multiple fruit categories, including apples, apricots, bananas, cherries, figs, grapes, kiwi fruit, peaches and nectarines, pears, plums and sloes, carob, and citrus fruits. Relative changes in production were quantified for two time windows: 1990–2022 and 2005–2022, the former corresponding to arbitrary selected broader time-window, the latter corresponding to our research period.

Temporal trends in fruit production were evaluated using ordinary least squares regression with year as the explanatory variable. To account for potential heteroscedasticity and temporal autocorrelation in time-series residuals, statistical inference was based on heteroscedasticity- and autocorrelation-consistent standard errors calculated using the Newey– West estimator (Newey and West, 1987). This approach is commonly used in time-series regression when residuals may show temporal dependence. Potential abrupt changes in fruit production dynamics were examined using the Pettitt test, a non-parametric test for detecting a single change point in a time series (Pettitt, 1979). The Pettitt test identifies the most likely year at which the median of a time series changes. As a robustness analysis, we also explored multiple structural breaks using the Bai–Perron multiple breakpoint procedure (Bai and Perron, 2003), implemented in the R package *strucchange* (Zeileis et al., 2002). This method allows detection of multiple structural changes in regression relationships and is widely used in economic and environmental time-series analyses. To assess whether structural changes occurred synchronously across different fruit production series, we examined the temporal distribution of detected breakpoints across all analysed crops. Clustering of breakpoints within specific time windows was tested using an exact binomial test, comparing the observed number of breakpoints in a given interval with the number expected under a random (uniform) temporal distribution. To test whether a common structural shift occurred across fruit production series, we fitted panel regression models with fruit-specific fixed effects. These models included fruit-specific intercepts and fruit-specific time trends, while testing for a shared post-break level shift and change in slope across all series. Statistical inference was based on cluster-robust standard errors clustered by fruit type to account for within-series dependence (Arellano, 1987).

All analyses were conducted in R version 4.5.2 (R Core Team, 2023). Log-linear population trend analyses were performed using *rtrim*, while generalized linear mixed models were fitted using *glmmTMB*. Structural break detection and time-series analyses were implemented using the packages *trend* (Pohlert, 2023), *strucchange* (Zeileis et al., 2002), *lmtest* (Zeileis and Hothorn, 2002), *sandwich* (Zeileis, 2004), *dplyr* (Wickham et al., 2023), and *ggplot2* (Wickham, 2016). Data manipulation and preparation were carried out using the *tidyverse* (Wickham et al., 2019).

## 3. Results

### 3.1. Population trends

Long-term population trend analysis using TRIM indicated a strong overall decline of fruit bat population over the monitoring period (−8.78% per year; 95% CI: −9.03 to −8.51; p < 0.001; Fig. 2). Between 2005 and 2022, the total number of fruit bats recorded across all monitored roosts declined from approximately 7200 individuals to approximately 1050 individuals. This corresponded to a decline in the population index from 100 to 14.6, equivalent to an estimated total decrease of 85.4% (95% CI: 84.5–86.4%). Inspection of annual indices and segment-specific analyses revealed three distinct temporal phases. During the first period (2005–2011), the population declined steeply, at −30.3% per year (95% CI: −31.0 to −29.7; p < 0.001). This was followed by a continued but less pronounced decline during 2012–2017 (−17.9% per year; 95% CI: −19.8 to −15.9; p < 0.001). In contrast, during 2018–2022 the trend reversed, and a significant increase was detected (+17.6% per year; 95% CI: 14.5–20.8; p = 0.007).

**Figure 2.**
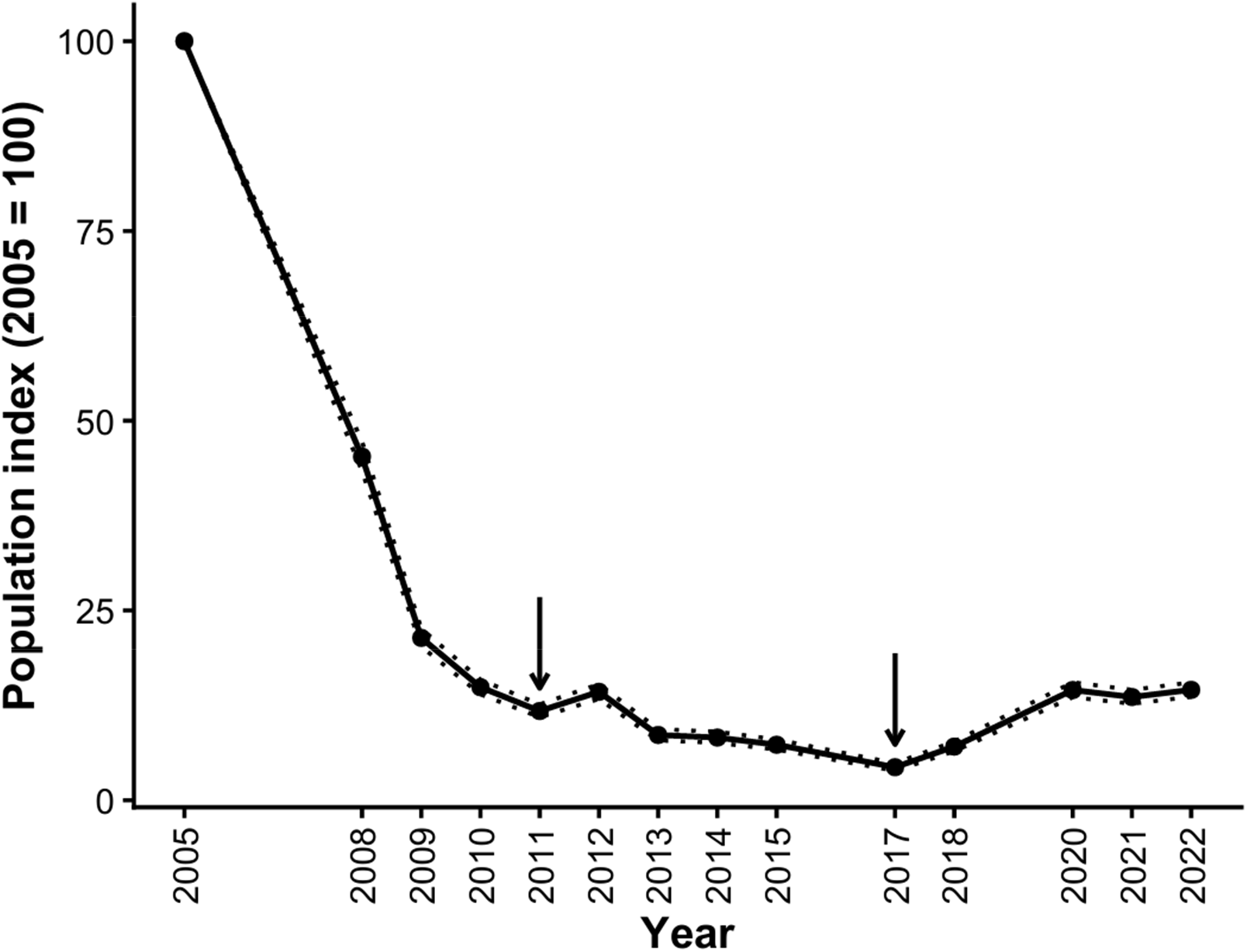
Temporal development of the population index Egyptian fruitbats between 2005 and 2022 based on monitoring of 21 colonies. Annual imputed indices were standardized to the baseline year 2005 (=100%). The grey ribbon represents 95% confidence intervals derived from model-based standard errors. Arrows indicate years (2011 and 2017) where significant year-to-year changes in the log-transformed population index were detected, identifying structural breakpoints subsequently used to define three analytical periods (2005– 2011, 2012–2017, 2018–2022) with different trends.

### 3.2. Effect of roost accessibility on population trends

Roosts in both accessibility categories showed significant declines in bat counts throughout the entire monitoring period; however, the rate of decline differed between them (Fig. 3). Colonies occupying easy-access roosts declined most steeply, by approximately −31.4% per year (95% CI: −38.4 to −23.6; p < 0.001). Colonies in hard-access roosts also declined significantly, but at a slower rate of −19.4% per year (95% CI: −24.5 to −13.9; p < 0.001). The difference between easy- and hard-access roosts was statistically significant (year × accessibility interaction: p = 0.011), indicating that roost accessibility influenced the rate of population decline. This interpretation was further supported by the fact that fruit bats disappeared completely from 87.5% of easily accessible roosts (N = 8), but from only 46.2% of hard-access roosts (N = 13).

**Figure 3.**
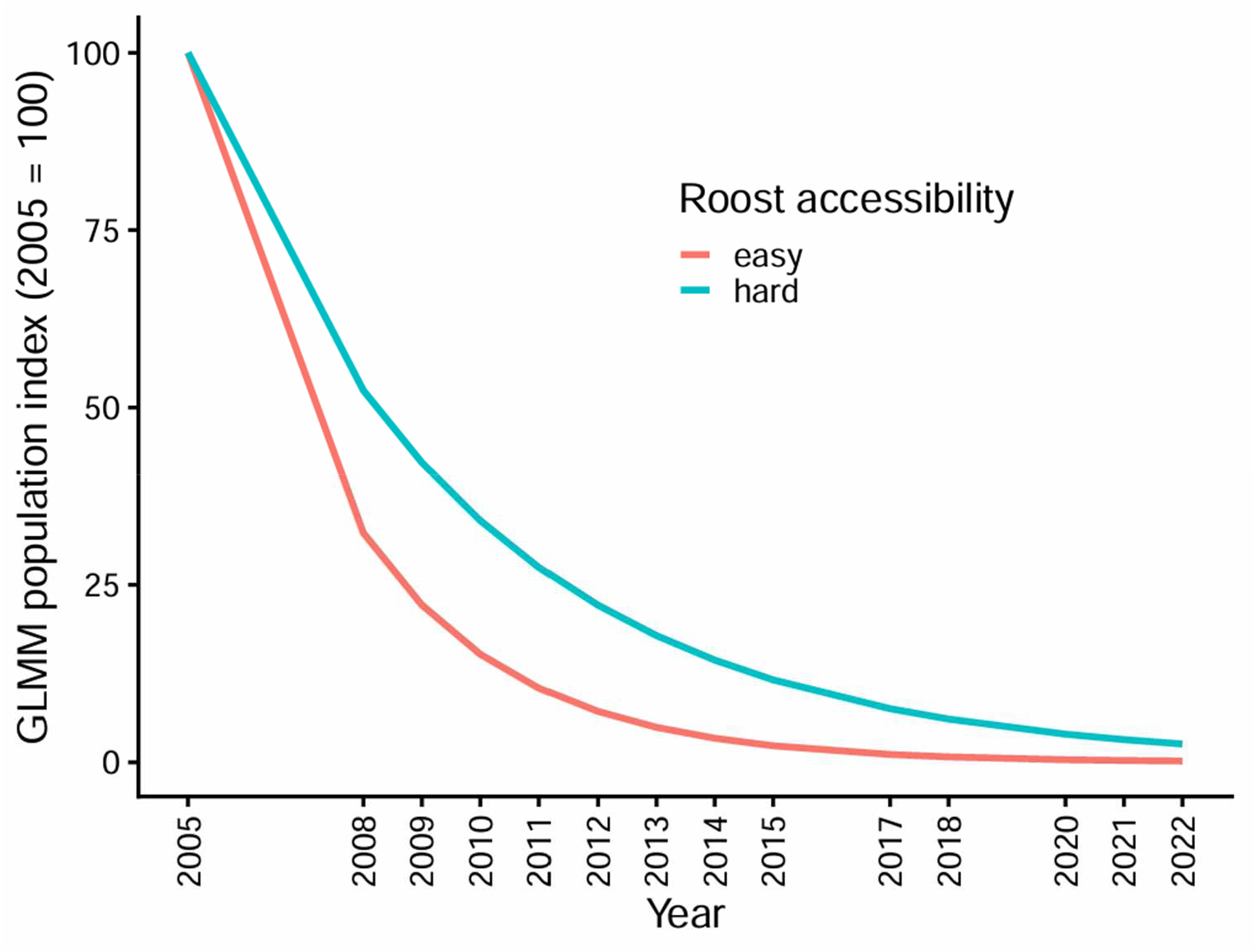
Modelled population trends of the Egyptian fruit bat (*Rousettus aegyptiacus*) in roosts differing in accessibility based on GLMM including year, roost accessibility, and their interaction, with site as a random effect. The population index was standardised to the first monitoring year (2005 = 100%). Colonies in easily accessible roosts declined more rapidly than those in less accessible sites.

### 3.3. Long-term trends in agricultural fruit production

Total fruit production showed a pronounced long-term decline in both analysed time windows (Fig. 4a). Between 1990 and 2022, aggregate fruit production decreased by 75.7%, whereas between 2005 and 2022 it declined by 58.8%. Across individual fruit types, production declined in most of the analysed series: nine fruit categories showed net decreases, whereas four showed overall increases over the longer time period (Appendix S1,Table S1). The largest declines between 1990 and 2022 were observed for grapes (−83.9%), citrus fruits (−73.5%), figs (−60.0%), and apples (−58.9%). By contrast, production increased in several fruit types, including kiwi fruit (+900%), plums and sloes (+82%), and peaches and nectarines (+70%). When focusing on the more recent period 2005–2022, which corresponds to the period of monitored fruit bat decline, production trends remained negative for most fruit types. Only plums and sloes (+119.3%) and carob (+7.7%) showed increases during this period, whereas all other crops declined (Appendix S1,Table S2).

**Figure 4.**
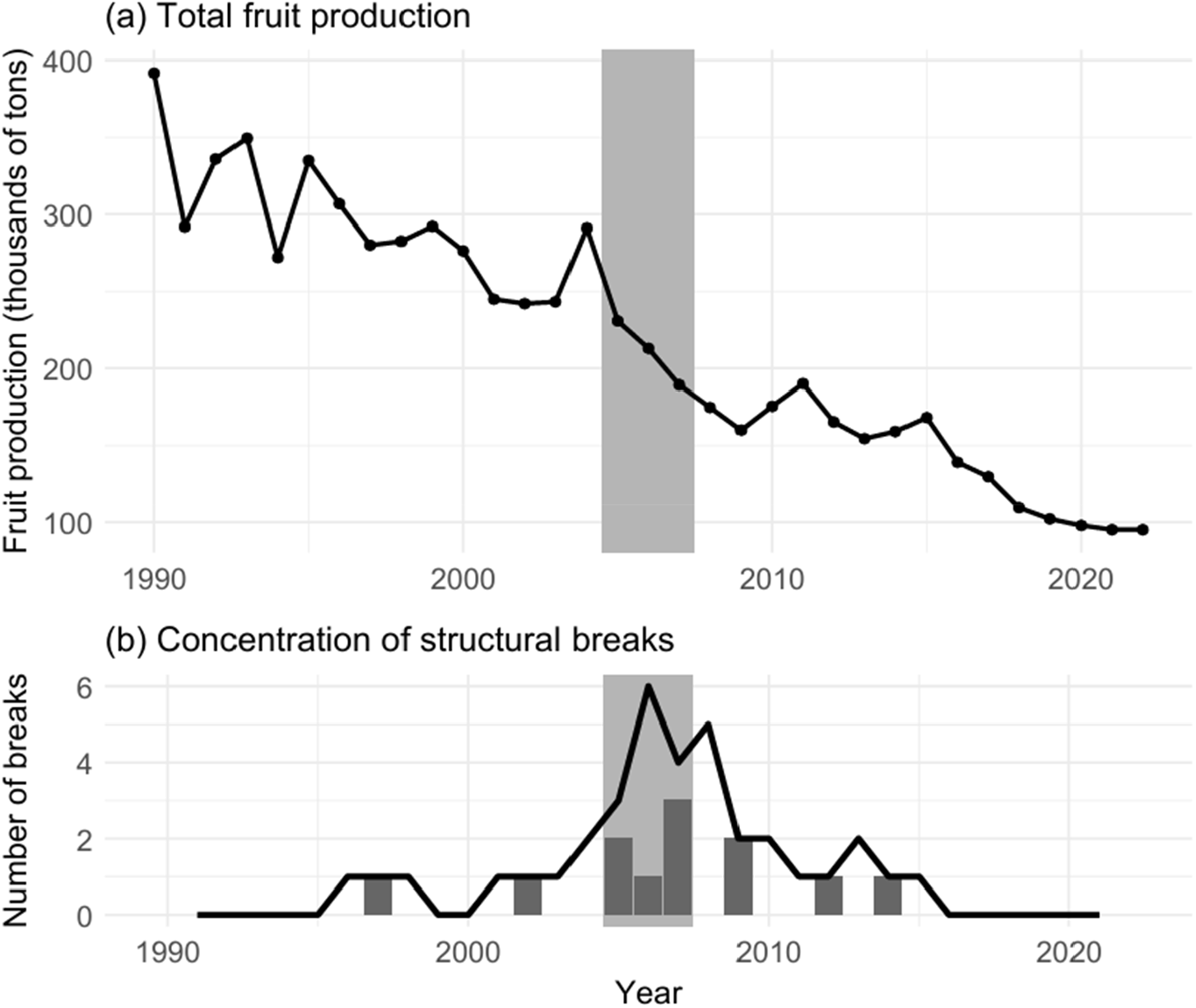
Annual total fruit production, expressed in thousands of tons, is presented in panel (a). The yearly number of statistically significant structural breaks detected in individual fruit production series using the Pettitt test is shown in panel (b). A centered three-year rolling sum of break occurrences, which smooths year-to-year variation and highlights periods of clustered structural change, is represented by the solid line The shaded interval marks the period 2005– 2007, during which the highest cross-series concentration of structural breaks occurred.

### 3.4. Structural changes in fruit production dynamics

Change-point analyses revealed a marked temporal clustering of structural breaks in fruit production series. The Pettitt change-point test detected statistically significant breakpoints in multiple crop time series, including cherries (U = 272, p < 0.0001), grapes (U = 272, p < 0.0001), apples (U = 270, p < 0.0001), citrus fruits (U = 258, p < 0.0001), apricots (U = 230, p < 0.001), and bananas (U = 216, p < 0.001). These breakpoints were concentrated mainly in the mid-2000s, suggesting a coordinated shift in production dynamics across several fruit crops.

When the temporal distribution of breakpoint years was examined across all crop series, the highest concentration of structural changes occurred between 2005 and 2007, when six breakpoints were detected across the analysed crops (Fig. 4b). A sliding-window analysis confirmed that this period represented the strongest clustering of breakpoints across the entire study period. Within the broader interval 2005–2011, which coincided with the steepest decline in the fruit bat population, eight of the 12 detected breakpoints occurred, whereas only 2.55 would be expected under a uniform temporal distribution. An exact binomial test confirmed that this concentration was highly unlikely to have arisen by chance (p < 0.001).

Panel regression analyses supported a coordinated shift in fruit production dynamics during the mid-2000s. In models allowing crop-specific trends, the estimated post-break slope change was negative for all candidate break years between 2004 and 2007, and statistically significant for 2004 (−0.0418, p < 0.05) and 2005 (−0.0365, p < 0.05). Although the post-break level shift was not significant in any specification, the joint Wald test supported a common structural change for 2005 (F = 3.11, p < 0.05), 2006 (F = 3.47, p < 0.05), and 2007 (F = 3.58, p < 0.05), with the 2004 model being marginally significant (F = 3.02, p = 0.050).

## 4. Discussion

### 4.1. Long-term population trends and roost accessibility as a proxy for roost disturbance

Between 2005 and 2022, the Egyptian fruit bat population on Cyprus declined by approximately 85% (mean annual decrease ∼9%), indicating a substantial population collapse. The decline proceeded in three phases: a steep initial drop (>30%/year, 2005–2011), a slower but continued decrease (∼18%/year, 2012–2017), and a partial recovery (2018–2022) suggesting that some driving factors may have weakened, though losses remain largely uncompensated. Our analysis also showed that population trends differed according to roost accessibility. Colonies occupying easily accessible roosts declined much more rapidly than colonies in less accessible roosts. This statistically supported difference suggests that roost accessibility may have played an important role in shaping population dynamics, most likely through variation in the intensity of human disturbance. Pteropodid bats worldwide are seriously threatened by direct persecution because they are often regarded as agricultural pests (Fujita and Tuttle, 1991). In the Egyptian fruit bat, such persecution has been documented in Israel, where eradication campaigns included fumigation of caves occupied not only by fruit bats but also by insectivorous bats (Makin and Mendelssohn, 1985). These actions caused severe declines in many insectivorous bat species, whereas fruit bats were reportedly less affected, probably because of their greater roosting plasticity and higher reproductive rate (Makin and Mendelssohn, 1985). In Cyprus, fruit bat control was historically organised by the Department of Agriculture and included direct killing, poisoning, roost fumigation, and the purchase of dead animals (Hadjisterkotis, 2006). Although this campaign officially ended in 1993 and all bat species are now legally protected, enforcement appears to have remained insufficient. For example, at least five cases of killing of fruit bats in roosts have reportedly appeared in Cypriot newspapers since 2006 (Patricia Radnor, ARC Kivotos, pers. comm.), and many additional cases may have gone unreported. We also found evidence of past eradication attempts, including burnt car tyres and empty shotgun cartridges, in 9 of 23 known fruit bat roosts, including two roosts designated as Natura 2000 sites (R. Lučan, pers. obs.).

At the same time, we doubt that direct persecution alone can account for the full magnitude of the decline documented here. During our study period, persecution focused on fruit bat roosts was almost certainly less intensive than the campaigns described for the 1970s by Spitzenberger (1979), who also reported rapid local disappearances from affected sites. Moreover, intensive persecution has also been reported from Israel, Lebanon, Turkey, and Egypt, yet no similarly dramatic long-term collapse has been documented there (Albayrak et al., 2008; Horáček et al., 2009). One factor that may help explain this contrast is the much more limited availability of suitable roosts on Cyprus. The island has relatively few caves, and most are small underground spaces extending only a few tens of metres at most (H. Nicolaou, pers. obs.). This contrasts strongly with mainland regions. For example, in Lebanon, where fruit bat abundance appeared stable or even increasing during the corresponding period, more than 500 larger caves providing potential alternative roosts have been registered (Karanouh, 2005). Under such conditions, the negative effects of human disturbance may be partly buffered because bats can respond by shifting to other roosts, including less accessible sites. On Cyprus, by contrast, fruit bats have very limited opportunities to relocate, making colonies particularly vulnerable to repeated disturbance.

Taken together, our results strongly suggest that roost disturbance contributed substantially to the collapse of the Cypriot fruit bat population. Although the available data do not allow direct quantification of disturbance intensity at individual roosts, the much steeper declines observed in easily accessible sites are fully consistent with this interpretation.

### 4.2. Declining availability of commercially produced fruits, its causes, and possible consequences

Our analyses of long-term agricultural production indicate a marked decline in the availability of commercially grown fruits that form an important part of the Egyptian fruit bat diet. In addition, structural break analyses showed that major changes in fruit production dynamics closely coincided with 2005–2007, the period during which the steepest decline in the fruit bat population was recorded. Although agricultural production is influenced by a range of socioeconomic factors, short-term and medium-term variation in Mediterranean agroecosystems is strongly shaped by weather and climate, especially by water availability. In such systems, rainfall amount and seasonal distribution are major determinants of productivity, and the effects of ongoing climate change are increasingly evident (Jacobsen et al., 2012; del Pozo et al., 2019).

Cyprus is especially vulnerable in this respect because of its limited water resources and strong dependence on climatic variability. Long-term reductions in rainfall, combined with rising temperatures, have repeatedly resulted in severe drought episodes (Markou et al., 2020; Cantonati et al., 2024). Of particular relevance is the fact that the most severe and prolonged drought period, including a year with the second-lowest rainfall recorded over more than a century, occurred between 2004 and 2008 (Michaelides and Pashiardis, 2008; Spinoni et al., 2015). This period coincides strikingly with both the major reorganisation of fruit production trajectories and the steepest phase of the fruit bat population collapse documented here. The drought also had major impacts on Cypriot agriculture and ecosystems more generally (Markou et al., 2020; Miltiadou et al., 2021). Taken together, these patterns suggest that the most severe phase of the fruit bat decline coincided with a climate-related reduction in food availability.

In addition to the decline in total fruit production, fruit availability to bats may have been further reduced by changes in orchard management. Across the Mediterranean region, including Cyprus, fruit production is becoming increasingly dependent on protective netting, a practice that was not typical of traditional orchard management (Adamides et al., 2022; Vuković et al., 2022). Various types of netting, including anti-hail, anti-bird, anti-bat, anti-insect, and photoselective nets, can reduce crop losses and contribute to more environmentally friendly agriculture by lowering pesticide use (Manja and Aoun, 2019). However, they also reduce or completely prevent access of fruit bats to cultivated fruit resources. This may impose particularly strong nutritional stress during extreme climatic events such as prolonged droughts and increasingly frequent heatwaves. As a result, the negative effect of declining commercially available fruit resources on fruit bats may have been even greater than implied by our production analyses alone.

No comparable long-term data are available for wild or ornamental plants, which also constitute an important part of the fruit bat diet (Korine et al., 1999; Del Vaglio et al., 2011). Nevertheless, it is highly likely that extreme climatic events, including prolonged drought and extreme heat, also affect this component of the diet. We suggest that the combined effects of drought and high temperatures may threaten fruit bats not only through reduced productivity of food plants, but also through changes in phenology. In Mediterranean ecosystems, increasing temperatures and more frequent heatwaves have advanced fruiting in many plant species, including important dietary resources for fruit bats (Gordo and Sanz, 2010). At the same time, spring-fruiting species tend to advance fruiting more strongly than autumn-fruiting species (Gordo and Sanz, 2005), potentially creating seasonal gaps in food availability. This may be particularly relevant because fruit bat food resources are already limited during summer and winter, whereas spring and autumn offer greater food availability (Herrera et al., 2008). Our previous work on the spatial activity and feeding ecology of Cypriot fruit bats suggested that as few as one or two major dietary items, notably *Ficus carica* and *Agave americana*, may be available in mid-summer on Cyprus (Lučan et al., 2016). The risk of temporary food bottlenecks is therefore likely to be high. Although fruit bats can consume leaves when preferred food is scarce (Kunz and Ingalls, 1994), folivory is unlikely to meet energetic demands over long periods or during energetically demanding stages such as reproduction (Korine et al., 2004). Last but not least it should be remembered that foraging strategy of fruit bat is largerly patterned by communal knowledge of foraging sites and kin-based site sharing, resulting in overall reduction of home ranges during time of poor food availability (Bachorec et al. 2020). A decline of food availability in traditional foraging sites might thus cause continuous starvation in considerable part of colony member, reduction of their capacity for searching unknown distant food resources and finally lead to a collapse of the colony as a vital unit of local population.

Water limitation may also have contributed directly to the decline. During the major drought of 2008, surface water reserves were nearly exhausted across the island (Michaelides and Pashiardis, 2008). Reduced water availability is known to affect bat reproduction strongly, as females under ecological stress may reduce or forego reproductive effort (Adams, 2010). Prolonged shortages of both food and water may therefore have increased mortality, reduced reproductive success, or both.

### 4.3. Fate of the missing population

An important question concerns the fate of the missing fraction of the Cypriot fruit bat population. Apart from individuals directly killed by people, the decline may reflect either increased mortality, for example due to starvation, or emigration from the island. We consider the latter scenario plausible for several reasons. First, throughout the study period, there has been no clear evidence of widespread mortality that could be attributed to exhaustion or starvation. Although bats generally have a cryptic lifestyle and often go unnoticed by the public, we would expect that conspicuous abnormal behaviour associated with severe nutritional stress, particularly in fruit bats, would occasionally be reported. Second, there is evidence of a recent expansion of Egyptian fruit bat populations within the thermo-Mediterranean zone of south-western Turkey and parts of Greece, including a westward shift of the species’ range margin along the southern Aegean coast of Anatolia (Benda et al., 2011; Strachinis et al., 2018).

Cyprus is separated from the Turkish coast by approximately 70 km of sea and from the Levantine coast by approximately 105 km. Previous studies indicate that the island was colonised from these mainland regions in the past (Hulva et al., 2012), and recent GPS-tracking data show that Egyptian fruit bats are capable of travelling distances exceeding 100 km within a single night (Amman et al., 2023). Movement from Cyprus to mainland areas is therefore physically feasible.

In addition, the Anatolian coastline is clearly visible at night from northern Cyprus, partly due to extensive artificial lighting associated with urbanised areas, which has increased markedly in recent decades and may facilitate navigation. Our field surveys along the southern Anatolian coast revealed highly favourable roosting and feeding conditions for fruit bats, including abundant populations occupying both natural roosts and synanthropic sites in urban environments such as Adana and Antalya (Benda et al., 2011).

Taken together, these observations suggest that emigration cannot be excluded as a contributing factor to the observed decline. However, this hypothesis remains untested and requires targeted investigation. A detailed population-level study, ideally combining genetic and movement data, is urgently needed to confirm or refute the role of emigration in shaping recent population dynamics.

### 4.4. Conservation implications

The unexpectedly severe decline of the only European population of this fruit bat represents a major conservation concern and calls for immediate, targeted action. At the same time, the slowing of the decline after 2012 and the positive trend observed in 2018–2022 suggest that at least some of the major stressors may have weakened, offering some hope for recovery if suitable conditions can be maintained or restored.

The most urgent priority is strict protection of all known roosts and effective prevention of human disturbance at remaining colony sites. Roost protection is a cornerstone of bat conservation because many species depend on a limited number of critical sites, making populations especially vulnerable to disturbance, exclusion, or roost loss (Neubaum et al., 2017; Frick et al., 2020; Browning et al., 2021). Several case studies further show that targeted roost protection and management can make substantial contributions to recovery and, in some cases, help reverse previous declines (Froidevaux et al., 2017; Wright et al., 2022).

In the longer term, conservation measures should also aim to ensure adequate access to food and water. The construction and maintenance of water sources for wildlife in arid environments is an established management practice (Rosenstock et al., 1999; Rich et al., 2019), but active support of food resources for frugivorous wildlife, and especially for bats, has been implemented much less frequently (Law et al., 2002). Because an important part of the diet of Cypriot fruit bats consists of fruits from plants commonly grown for non-commercial purposes, such as mulberry (*Morus* spp.) and loquat (*Eriobotrya japonica*) (Del Vaglio et al., 2011), conservation management should also consider promoting the planting of these species where appropriate. Effective management may even require consideration of more controversial measures, such as the controlled use of non-native food plants, for example Persian lilac (*Melia azedarach*) or century plant (*Agave americana*), if these provide critical food during nutritionally limiting periods and have high dietary value for fruit bats (Lučan et al., 2016).

Finally, the Egyptian fruit bat has long been perceived as an agricultural pest, and this negative perception may still persist in some communities. In addition, complaints submitted to Cypriot conservation authorities often concern the conspicuous faecal marks left by fruit bats on house facades (H. Nicolaou, unpublished data), which can create public nuisance issues. Notably, such signs were also used by researchers to confirm the presence of fruit bats in localities surveyed on Cyprus (Benda et al., 2007, 2018). Effective conservation of this species will therefore require not only habitat and roost protection, but also targeted public outreach, conflict mitigation, and practical measures aimed at reducing negative interactions between fruit bats and people.

## 5. Conclusion

Our study documents a severe collapse of the Cypriot population of the Egyptian fruit bat, with an estimated decline of more than 85% between 2005 and 2022. The decline was most pronounced during the early phase of monitoring, subsequently slowed, and was followed by signs of partial recovery in recent years. Multiple lines of evidence indicate that this collapse was most likely driven by the combined effects of roost disturbance and reduced food availability associated with climate-related environmental change. Colonies in easily accessible roosts declined more rapidly than those in less accessible sites, supporting an important role of human disturbance, while long-term declines and structural shifts in fruit production coincided with the most severe phase of population decline.

At the same time, the fate of the missing portion of the population remains uncertain. Although large-scale mortality cannot be ruled out, there was no clear evidence of widespread starvation-related mortality. Emigration from Cyprus to nearby mainland regions is physically possible and is supported indirectly by recent range expansion in neighbouring areas and favourable ecological conditions along the southern Anatolian coast. However, this hypothesis remains unresolved and requires targeted investigation.

Given that this population represents the only insular and genetically distinctive population of the Egyptian fruit bat in Europe, its decline is of considerable conservation concern. Immediate protection of roost sites, combined with measures to improve access to food and water resources, should be considered a priority. At the same time, future research should focus on resolving the relative roles of mortality and emigration in driving population change, as this distinction has important implications for effective conservation management.

## CRediT authorship contribution statement

**Radek K. Lučan**: Writing – review & editing, Writing – original draft, Visualization, Methodology, Investigation, Data analysis, Data curation, Conceptualization. **Haris Nicolaou**: Writing – review & editing, Investigation, Data curation. **Tomáš Bartonička**: Writing – review & editing, Investigation. **Eric Bachorec**: Writing – review & editing, Data analysis. **Martin Šálek**: Writing – review & editing, Investigation. **Šimon Řeřucha**: Writing – review & editing, Investigation. **Petr Jedlička**: Writing – review & editing, Investigation. **Ivan Horáček**: Writing – review & editing, Supervision, Methodology, Investigation, Funding acquisition, Conceptualization.

## Acknowledgement

We would like to thank to Petr Benda, Elena Erotokritou, Pavel Hulva, Helena Jahelková, George Konstantinou, Magdalena Lučanová and Anna Lučanová for their help with fieldwork and we thank Tereza Stříbrná for providing population genetic data on fruit bats.The project was supported by the Grant Agency of Academy of Sciences of the Czech Republic (project no. IAA601110905) and the Grant Agency of the Czech Republic (project no. 206/09/0888).

## Data availability

Data and complete analytical workflows with R scripts will be made available on request.

## Notes

### Competing Interest Statement

The authors have declared no competing interest.

