## Appendix S1 for "Population collapse of the Egyptian fruit bat on Cyprus (2005−2022) was likely driven by roost disturbance and declining food availability linked to climate change"

Supplementary information

| **Table S1. Linear trend estimates for annual fruit production series (1990–2022)** | | | | | | |
| --- | --- | --- | --- | --- | --- | --- |
| *Estimates are based on ordinary least squares regressions of annual production on year. Statistical inference is based on Newey–West HAC standard errors.* | | | | | | |
| *HAC SE = heteroscedasticity- and autocorrelation-consistent standard error.* | | | | | | |
| **Fruit** | **Slope per year** | **HAC SE** | **t value** | **p value** | **95% CI low** | **95% CI high** |
| **Total fruit production** | **-8291.379** | **259.651** | **-31.933** | **P < 0.001** | **-8800.296** | **-7782.463** |
| All citruses | -3530.862 | 350.058 | -10.087 | **P < 0.001** | -4216.975 | -2844.749 |
| Grapes | -4093.393 | 433.354 | -9.446 | **P < 0.001** | -4942.767 | -3244.020 |
| Cherries | -35.337 | 5.787 | -6.106 | **P < 0.001** | -46.680 | -23.994 |
| Apples | -262.882 | 51.336 | -5.121 | **P < 0.001** | -363.501 | -162.263 |
| Pears | -22.695 | 5.014 | -4.527 | **P < 0.001** | -32.522 | -12.869 |
| Apricots | -38.960 | 11.892 | -3.276 | **P < 0.01** | -62.269 | -15.651 |
| Bananas | -136.817 | 48.825 | -2.802 | **P < 0.01** | -232.513 | -41.120 |
| Kiwi fruit | -6.874 | 4.159 | -1.653 | 0.108 | -15.024 | 1.277 |
| Figs | -46.028 | 28.225 | -1.631 | 0.113 | -101.350 | 9.294 |
| Plums and sloes | 10.454 | 8.401 | 1.244 | 0.223 | -6.011 | 26.919 |
| Peaches and nectarines | 16.906 | 29.027 | 0.582 | 0.564 | -39.987 | 73.800 |
| Carob | 33.337 | 63.755 | 0.523 | 0.605 | -91.622 | 158.297 |

| **Table S2. Linear trend estimates for annual fruit production series (2005–2022)** | | | | | | |
| --- | --- | --- | --- | --- | --- | --- |
| *Estimates are based on ordinary least squares regressions of annual production on year. Statistical inference is based on Newey–West HAC standard errors.* | | | | | | |
| *HAC SE = heteroscedasticity- and autocorrelation-consistent standard error.* | | | | | | |
| **Fruit type** | **Slope per year** | **HAC SE** | **t value** | **p value** | **95% CI low** | **95% CI high** |
| **Total fruit production** | **-7300.157** | **624.169** | **-11.7** | **P < 0.001** | **-8523.529** | **-6076.785** |
| All citruses | -4874.053 | 535.41 | -9.1 | **P < 0.001** | -5923.457 | -3824.649 |
| Apples | -477.183 | 64.935 | -7.35 | **P < 0.001** | -604.456 | -349.909 |
| Figs | -163.8 | 24.587 | -6.66 | **P < 0.001** | -211.99 | -115.61 |
| Peaches and nectarines | -138.557 | 22.541 | -6.15 | **P < 0.001** | -182.738 | -94.376 |
| Plums and sloes | 61.099 | 11.105 | 5.502 | **P < 0.001** | 39.334 | 82.864 |
| Pears | -55.347 | 10.148 | -5.45 | **P < 0.001** | -75.236 | -35.457 |
| Cherries | -12.557 | 2.735 | -4.59 | **P < 0.001** | -17.918 | -7.196 |
| Bananas | -64.99 | 17.425 | -3.73 | **P < 0.01** | -99.143 | -30.836 |
| Grapes | -1112.157 | 437.92 | -2.54 | **P < 0.05** | -1970.48 | -253.834 |
| Kiwi fruit | -2.221 | 0.93 | -2.39 | **P < 0.05** | -4.045 | -0.397 |
| Apricots | -37.632 | 19.205 | -1.96 | 0.0677 | -75.273 | 0.009 |
| Carob | 211.676 | 145.445 | 1.455 | 0.174 | -73.397 | 496.749 |
